# Inhibition of the Receptor for Advanced Glycation End-Products in Acute Respiratory Distress Syndrome: A Randomised Laboratory Trial in Piglets

**DOI:** 10.1101/405423

**Authors:** Jules Audard, Thomas Godet, Raiko Blondonnet, Jean-Baptiste Joffredo, Bertille Paquette, Corinne Belville, Marilyne Lavergne, Christelle Gross, Justine Pasteur, Damien Bouvier, Loic Blanchon, Vincent Sapin, Bruno Pereira, Jean-Michel Constantin, Matthieu Jabaudon

**Affiliations:** Department of Perioperative Medicine, CHU Clermont-Ferrand, Clermont-Ferrand, France; Université Clermont Auvergne, CNRS UMR 6293, INSERM U1103, GReD, Clermont-Ferrand, France; Department of Dermatology, CHU Clermont-Ferrand, Clermont-Ferrand, France; Department of Medical Biochemistry and Molecular Biology, CHU Clermont-Ferrand, Clermont-Ferrand, France; Biostatistical and Data Management Unit, Department of Clinical Research and Innovation (DRCI), CHU Clermont-Ferrand, Clermont-Ferrand, France

**Keywords:** acute respiratory distress syndrome, receptor for advanced glycation end-products, animal model, alveolar fluid clearance, therapy

## Abstract

**Background:** The receptor for advanced glycation end products (RAGE) modulates the pathogenesis of acute respiratory distress syndrome (ARDS). RAGE inhibition was recently associated with attenuated lung injury and restored alveolar fluid clearance (AFC) in a mouse model of ARDS. However, clinical translation will first require assessment of this strategy in larger animals.

**Methods:** Forty-eight anaesthetised Landrace piglets were randomised into a control group and three treatment groups. Animals allocated to treatment groups underwent orotracheal instillation of hydrochloric acid i) alone; ii) in combination with intravenous administration of a RAGE antagonist peptide (RAP), a S100P-derived peptide that prevents activation of RAGE by its ligands, or iii) in combination with intravenous administration of recombinant soluble (s)RAGE that acted as a decoy receptor. The primary outcome measure was net AFC at 4 h. Arterial oxygenation was assessed hourly for 4 h and alveolar-capillary permeability, alveolar inflammation, lung histology and lung mRNA expression of the epithelial sodium channel (α1-ENaC), α1-Na,K-ATPase and aquaporin (AQP)-5 were assessed at 4 h.

**Findings:** Treatment with either RAP or sRAGE improved net AFC rates (median [interquartile range], 21.2 [18.8–21.7] and 19.5 [17.1–21.5] %/h, respectively, versus 12.6 [3.2–18.8] %/h in injured, untreated controls), improved oxygenation and decreased alveolar inflammation and histological evidence of tissue injury after acid-induced ARDS. RAGE inhibition also restored lung mRNA expression of α1-Na,K-ATPase and AQP-5.

**Interpretation:** RAGE inhibition restored AFC and attenuated lung injury in a piglet model of acid-induced ARDS.

**Funding:** Auvergne Regional Council, *Agence Nationale de la Recherche, Direction Générale de l’Offre de Soins*.

**Research in Context:** *Evidence before this study:* The acute respiratory distress syndrome (ARDS), a clinical syndrome of diffuse pulmonary oedema and inflammation, currently lacks effective therapies and is associated with high mortality and morbidity. The degrees of lung epithelial injury and of alveolar fluid clearance (AFC) impairment, as evaluated by plasma levels of soluble receptor for glycation end-products (RAGE), are major prognostic factors in ARDS and potential therapeutic targets for ongoing research. For example, targeting RAGE with recombinant sRAGE or an anti-RAGE monoclonal antibody has proven beneficial in a translational mouse model of acid-induced ARDS.

*Added value of this study:* In a piglet model of acid-induced ARDS, treatment with RAGE antagonist peptide or recombinant sRAGE restored AFC and attenuated the features of lung injury, thereby confirming, in the closest evolutionary model species to humans, previous evidence from rodent models that modulation of RAGE may be a therapeutic option for ARDS. Although this is an important step towards future clinical translation, future studies should assess the best methods to modulate RAGE and further confirm the safety of manipulating this pathway in patients with ARDS.

## INTRODUCTION

Acute respiratory distress syndrome (ARDS) is a frequent cause of respiratory failure and death in critically ill patients [1]. ARDS currently lacks effective therapies, although lung-protective ventilation and reasoned fluid management remain essential for better outcomes [2,3]. ARDS is a clinical syndrome [4] characterised by diffuse alveolar epithelial and lung endothelial injury that leads to increased permeability pulmonary oedema, alveolar filling and respiratory failure [2]. The resolution of alveolar oedema and ARDS requires that the alveolar epithelial fluid transport function remain intact, which suggests that a strategy aimed at improving alveolar fluid clearance (AFC) may be beneficial for recovery from ARDS [5,6]. The main mechanism responsible for the reabsorption of the water fraction of the oedema fluid from the airspaces of the lungs is active ion transport across the alveolar epithelium, which occurs primarily through the operation of the epithelial sodium channel (ENaC), Na,K-ATPase and aquaporin (AQP)-5 [7].

The lung alveolar type (AT)-1 cells abundantly express the receptor for advanced glycation end-products (RAGE) as a transmembrane pattern-recognition receptor [8,9], suggesting that RAGE may play a central role in the pathogenesis of ARDS [8,10–13]. The activation of RAGE modulates cell signalling that leads to a sustained inflammatory response through various intracellular signalling pathways that typically lead to pro-inflammatory activation of nuclear transcription factor NF-κB and upregulation of RAGE itself [9]. Plasma levels of soluble RAGE (sRAGE) show a positive association with the extent and severity of lung injury, the degree of AFC impairment and worse clinical outcomes in ARDS [10–12,14–16].

Recent findings in genetically deficient (RAGE^-/-^) mice or rats treated with an anti-RAGE antibody suggest that targeting RAGE might be a beneficial therapy for treatment of experimental sepsis and pneumonia [17,18]. In addition, the administration of sRAGE, which acts as a decoy receptor to prevent interactions between transmembrane RAGE and its ligands, decreased the alveolar inflammation and lung permeability in mice with lipopolysaccharide (LPS)-induced lung injury [19]. Our team recently showed that treatment with either an anti-RAGE monoclonal antibody or sRAGE attenuated lung injury, improved arterial oxygenation and decreased alveolar inflammation in a translational model of acid-injured mice [20]. The anti-RAGE therapies restored AFC and increased lung expression of AQP-5 in alveolar cells, thereby providing a possible link between RAGE modulation and the mechanisms of lung epithelial injury and repair relevant to clinical ARDS [7].

However, this type of strategy for RAGE inhibition has never been tested in a large animal model, either to determine the precise functional and biological effects of RAGE modulation in ARDS or in preparation for translation of this strategy to the clinical setting as an ARDS treatment. The aim of the present randomised trial was to investigate the potential therapeutic roles of either recombinant sRAGE or a RAGE antagonist peptide (RAP) on AFC and to establish the main features of experimental ARDS in a piglet model of hydrochloric acid (HCl)-induced ARDS. RAP is a small S100P-derived peptide with similar blocking effects to those of anti-RAGE monoclonal antibody, as it prevents RAGE from binding with several of its most important ligands, including HMGB1 and S100 proteins, in cancer cells in vitro and in vivo [21]. We also explored whether RAGE inhibition influenced lung expression of the alveolar epithelial ENaC channel, Na,K-ATPase and AQP-5.

## MATERIALS and METHODS

### Animal model

Animals were maintained and all procedures were performed with the approval of the ethics committee of the French *Ministère de l’Education Nationale, de l’Enseignement Supérieur et de la Recherche* in the *Centre International de Chirurgie Endoscopique*, School of Medicine - University of Clermont-Ferrand (approval number 01505.03). All experiments were performed in accordance with the “Animal Research: Reporting In Vivo Experiments” (ARRIVE) guidelines (**Supplementary Checklist**) [22].

Two-month-old white Landrace male piglets with mean (± standard deviation (SD)) weights of 10.1 (± 1.1) kg were restricted from food overnight but allowed free access to water, before receiving premedication with intramuscular azaperone (2 mg.kg^-1^). General anaesthesia was then induced with intravenous propofol (3 mg.kg^-1^) and sufentanil (0.3 µg.kg^-1^) prior to orotracheal intubation (6-mm ID cuffed endotracheal tube), and anaesthesia was maintained with continuous intravenous infusion of propofol (5 mg.kg^-1^.h^-1^) and remifentanil (10-20 µg.kg^-1^.h^-1^). The body temperature of the pigs was kept at approximately 38°C using warm blankets (Medi-therm II, Gaymar Industries, Orchard Park, NY, USA). Mechanical ventilation was delivered, with the pigs in the supine position, using volume-controlled ventilation, a tidal volume of 6 ml.kg^-1^, a positive end-expiratory pressure (PEEP) of 5 cmH_2_O and an oxygen inspired fraction (FiO_2_) of 40% (Engström Carestation, GE Healthcare, Chicago, IL, USA). The respiratory rate was adjusted to maintain the end-tidal carbon dioxide between 35 and 45 mmHg. Central venous access through the jugular vein and catheterisation of the femoral artery allowed retrieval of serial blood samples and continuous hemodynamic monitoring (arterial pressure, cardiac index and extravascular lung water (EVLW), as indexed to body weight [23]) with a PiCCO+ device (Maquet, Rastatt, Germany). The electrocardiogram activity and the peripheral oxygen saturation (SpO_2_) arterial pressure were also monitored continuously (IntelliVue MP40, Philips, Amsterdam, The Netherlands).

A total of 48 piglets was randomly allocated to four groups by means of computer software (Microsoft Office Excel 2003, Microsoft Corporation, Redmond, WA, USA). The “Sham” group was composed of control animals without lung injury (n=12). The “HCl group” consisted of animals with HCl-induced lung injury (n=12). Animals with HCl-induced lung injury and receiving intravenous treatment with RAP (EMD Millipore, Burlington, MA, USA) (3 µg.kg^-1^) defined the “RAP group” (n=12). The “sRAGE group” (n=12) included animals with HCl-induced lung injury that also received intravenous treatment with sRAGE (3 mg.kg^-1^) (RAGE (N-16) peptide, Santa Cruz Biotechnology, Dallas, TX, USA). Intravenous RAP or sRAGE was administered 30 minutes prior to the HCl instillation, based on findings from previous studies [17,20].

Acid aspiration–induced ARDS was produced by intratracheal instillation of 0.05 M HCl, pH 1.41 (4 ml.kg^-1^ body weight), over 3 min at the level of the carina [24]. Based on previous studies, lung injury was considered established when the PaO_2_/FiO_2_ ratio decreased to 25% from the baseline, approximately one hour after airway HCl instillation [24,25].

Animals were maintained under anaesthesia and mechanical ventilation for four hours after HCl instillation. At the end of ventilation, and after arterial blood sampling and a lung bronchoalveolar lavage (BAL) with 50 mL of saline, the piglets were sacrificed with intravenous pentobarbital (150 mg.kg^-1^).

### Outcome measures

#### Primary outcome

The primary outcome was the net AFC rate. Undiluted pulmonary oedema fluid samples were collected from the animals at baseline and four hours later, as previously described [12,26–31]. Briefly, a soft 14-Fr-gauge suction catheter (ConvaTec, Lejre, Denmark) was advanced into a wedged position in a distal bronchus via the endotracheal tube and oedema fluid was collected in a suction trap by applying gentle suction. All samples were centrifuged at 240 × g at 4°C for 10 min in a refrigerated centrifuge. The supernatants were collected and the total protein concentration was determined in duplicate with a colorimetric method (Pierce BCA Protein Assay Kit, ThermoFisher Scientific, Waltham, MA, USA). Because the rate of clearance of oedema fluid from the alveolar space is much faster than the rate of protein removal [32], the net AFC rate was calculated as Percent AFC = 100 × [1 - (initial oedema protein/final oedema total protein)] and thereafter was reported as %/h. All samples had a coefficient of variation of less than 10%.

#### Secondary outcomes

Secondary outcomes were major criteria for experimental ARDS, as recommended by the *American Thoracic Society* [33].

At baseline and every hour for four hours, arterial blood gases were measured to assess PaO_2_/FiO_2_, PaCO_2_, pH and serum lactate (Epoc^^®^^ Blood Analysis System, Siemens Healthineers, Erlangen, Germany), and respiratory (tidal volume, inspiratory plateau pressure, compliance of the respiratory system, driving pressure) and hemodynamic (mean arterial pressure, cardiac index, EVLW) parameters were collected.

In addition to measuring the ELVW through transpulmonary thermodilution, the alteration of the alveolar-capillary barrier was assessed by measuring the BAL level of total protein at four hours as a surrogate for alveolar oedema.

Alveolar inflammation was assessed by duplicate determination of the levels of proinflammatory cytokines, including tumour necrosis factor (TNF)-α, interleukin (IL)-6, IL-1β and IL-18, in the BAL at four hours. These determinations were made using the Bio-Plex 200 System, which is based on Luminex xMAP Technology (Bio-Rad, Hercules, CA, USA) and a Milliplex MAP Kit (Luminex xMAP technology, EMD Millipore, Burlington, MA, USA). All samples had a coefficient of variation of less than 10%.

After sacrificing the piglets, whole lungs were removed, fixed with alcoholic acetified formalin and embedded in paraffin. Slices at 10-µm thickness were stained with hematoxylin and eosin (Sigma-Aldrich, Saint-Louis, MO, USA). Histological evidence of lung injury was assessed using a standardised, validated histology injury score derived from the following calculation: score = [20×(i) + 14×(ii) + 7×(iii) + 7×(iv) + 2×(v)] / (number of fields × 100) (**Supplementary Table 1**) [33].

In parallel, total RNA was isolated from the lung with an RNA extraction kit (RNeasy^®^ Mini Kit, Qiagen, Hilden, Germany). The gene expression levels of α1-ENaC (NCBI Reference Sequence, NM_213758.2), α1-Na,K-ATPase (NM_214249.1), AQP-5 (NM_001110424.1) and 36B4 (a housekeeping gene) were assessed using the quantitative real-time reverse transcription polymerase chain reaction. The aim was to explore potential mechanisms related to the lung epithelial channels involved in transepithelial fluid transport through which anti-RAGE therapies might restore AFC [20]. Threshold levels of mRNA expression (ΔΔCt) were normalised to the housekeeping gene. The values represent the mean of triplicate samples ± SD and the data are representative of three independent observations. The primers are listed in **Supplementary Table 2**.

The researchers performing animal experiments and collecting samples were not blinded to group allocation. However, the researchers who performed measurements from the biological samples (e.g., blood gas analyses, protein and cytokine measurement, histology score) and the statistician who performed analyses were blinded.

### Statistical Analysis

All analyses were performed using Prism 6 (GraphPad Software, La Jolla, CA) or Stata version 14 (StataCorp, College Station, TX).

Categorical data were expressed as numbers and percentages, and quantitative data as mean and SD or median and interquartile range [IQR] according to their statistical distribution. Baseline characteristics between groups were compared using Student’s t-test or Mann–Whitney test were considered for quantitative parameters according to the t-test assumption (normality assumption using Shapiro–Wilk test and homoscedasticity with Fisher–Snedecor test). Categorical data were compared among groups using the Chi-square test or Fisher’s exact test.

Analysis of physiological parameters with repeated measurements was carried out by two-way repeated measures analyses of variance (ANOVA) when appropriate; Kruskal-Wallis test with Bonferroni tests were used for pairwise comparisons. Time × group interactions and post-hoc comparisons were verified using random effects models to analyse longitudinal evolution of variables: (i) considering between-and within-subject variability (random subject effects: random intercept and slope) and (ii) evaluating fixed effects: group, time and time × group interaction. The residual normality was checked for all models. Values were log-transformed for all variables to achieve normality prior to performing random effects models.

According to the principles of the 3Rs (Replacement, Reduction and Refinement), the smallest number of animals was used (12 animals in each group) that could detect a mean difference of 5%/h (SD = 4) in the net AFC rate at four hours after lung injury (primary outcome) between HCl-injured animals and HCl-injured animals receiving sRAGE or RAP, while still considering the alpha and beta risks of 5% (bilateral) and 10%, respectively. A statistical power of 90% was considered sufficient to allow multiple comparisons between groups.

## RESULTS

### Alveolar Fluid Clearance

A significant between-group difference was detected in net AFC rates measured after four hours of mechanical ventilation (P = 0.02) (**Fig. 1**). The net AFC rate was significantly decreased in HCl-injured piglets (12.6 [3.2–18.8] %/h) when compared with sham animals (17.9 [14.4–25.5] %/h). By contrast, treatment with RAP (21.2 [18.8–21.7] %/h) or sRAGE (19.5 [17.1–21.5] %/h) restored AFC in the HCl-injured animals.

**Figure 1.**
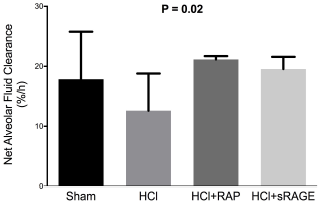
RAGE inhibition restores alveolar fluid clearance after acid-induced lung injury. Measurement of net alveolar fluid clearance (AFC) rate as a marker of epithelial function in uninjured (Sham), acid-injured (HCl) and acid-injured piglets treated with RAGE antagonist peptide (HCl+RAP) or recombinant sRAGE (HCl+sRAGE) (n = 12 per group at each time point). Values are reported as medians and interquartile ranges.

### Arterial Oxygenation

Two-way repeated-measurement ANOVA indicated a group effect (P = 0.001), a time effect (P = 0.03) and a significant interaction (P <10^-4^) with a detrimental effect of HCl-induced ARDS on the course of PaO_2_/FiO_2_, when compared with the absence of ARDS or with the use of an anti-RAGE therapy with either RAP or sRAGE in animals with ARDS (**Fig. 2**). Arterial oxygenation, as assessed by PaO_2_/FiO_2_, decreased by 25% at one hour (301 [223–405] mmHg) in untreated ARDS animals, and this decrease remained stable throughout the experiment in those animals (314 [186–447] mmHg at four hours). In both groups of ARDS animals treated with RAP or sRAGE, arterial oxygenation was preserved at all time points when compared with otherwise untreated, injured animals (419 [385–452] and 431 [412–466] mmHg at four hours, respectively).

**Figure 2.**
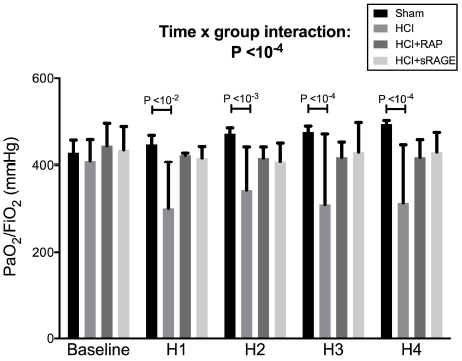
RAGE inhibition improves arterial oxygenation after acid-induced lung injury. Arterial oxygen tension (PaO_2_)/inspiratory oxygen fraction (FiO_2_) in uninjured (Sham), acid-injured (HCl) and acid-injured piglets treated with RAGE antagonist peptide (HCl+RAP) or recombinant sRAGE (HCl+sRAGE) (n = 12 per group at each time point). Values are reported as medians and interquartile ranges.

### Alteration of the Alveolar-Capillary Barrier

A significant between-group difference was noted in BAL levels of total protein measured after four hours of mechanical ventilation (P = 10^-4^) (**Fig. 3A**). HCl-induced ARDS was associated with increased BAL protein (6.1 [3.1–7.7] g.L^-1^), when compared with sham animals (0.3 [0.2–0.7] g.L^-1^) and animals treated with either RAP (1.2 [0.9–2.1] g.L^-1^) or sRAGE (1.3 [1.0–2.1] g.L^-1^).

**Figure 3.**
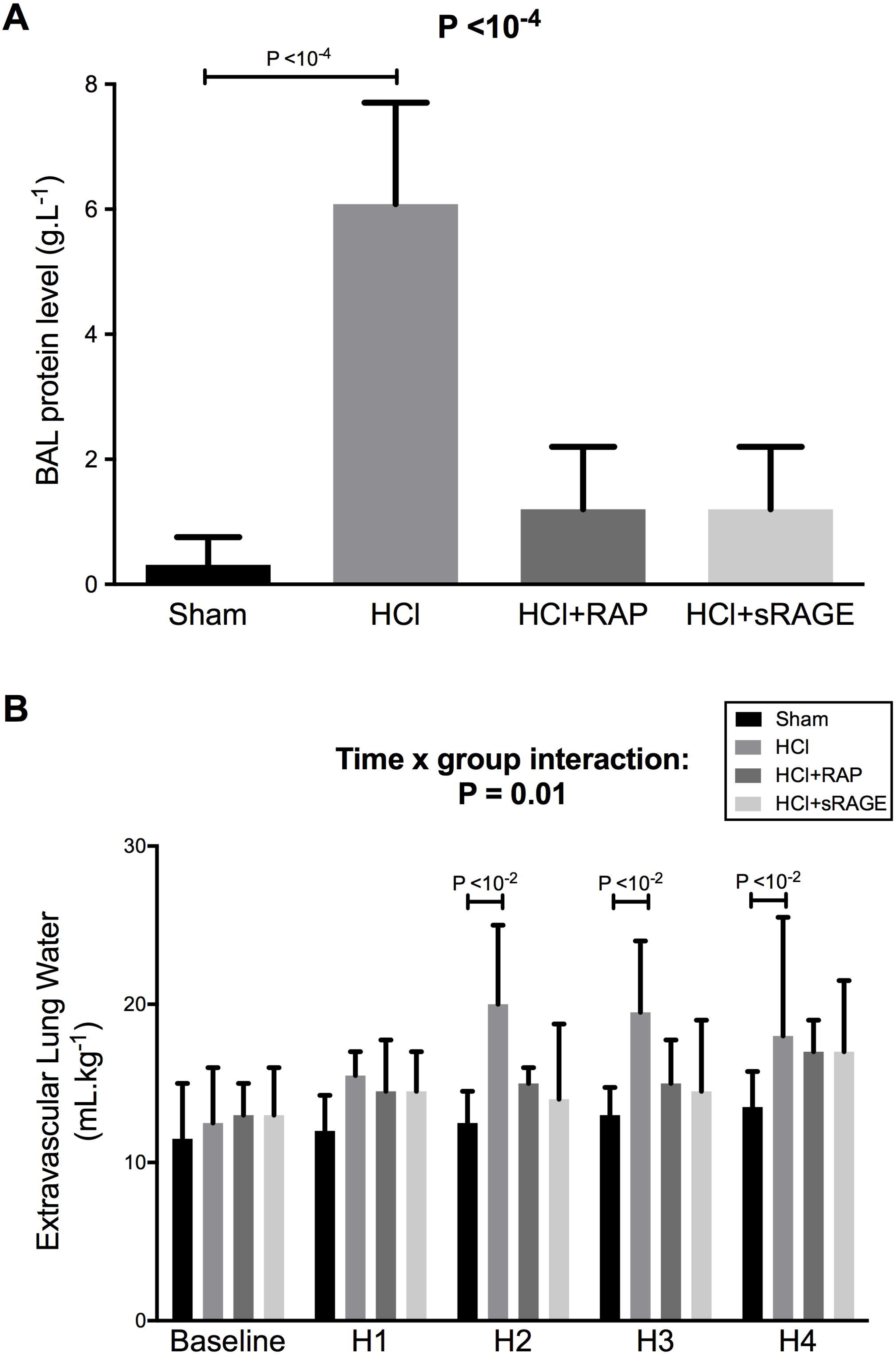
RAGE inhibition decreases alveolar-capillary permeability after acid-induced lung injury. (**A**) Level of total protein in the bronchoalveolar lavage (BAL) fluid from uninjured (Sham), acid-injured (HCl) and acid-injured piglets treated with RAGE antagonist peptide (HCl+RAP) or recombinant sRAGE (HCl+sRAGE) (n = 12 per group). (**B**) Extravascular lung water, as measured by transpulmonary thermodilution (Picco+, Pulsion SA) and indexed to body weight, in uninjured (Sham), acid-injured (HCl) and acid-injured piglets treated with RAGE antagonist peptide (HCl+RAP) or recombinant sRAGE (HCl+sRAGE) (n = 12 per group at each time point). Values are reported as medians and interquartile ranges.

Two-way repeated-measurement ANOVA indicated a group effect (P = 0.03), a time effect (P <10^-4^), and a significant interaction (P = 0.01) with an incremental effect of HCl-induced injury on the course of EVLW, when compared with the absence of injury or with the use of RAP or sRAGE (**Fig. 3B**). EVLW increased significantly at two, three and four hours of mechanical ventilation (P <0.01 at all time points) in HCl-injured animals (20.5 [15.0–25.0] mL.kg^-1^ at four hours) as compared with sham animals (14.7 [12.0–15.5] mL.kg^-1^ at four hours). However, EVLW was lower in animals treated with RAP or sRAGE (16.8 [14.0–19.0] mL.kg^-1^ and 17.6 [14.5–21.0] mL.kg^-1^ at four hours, respectively) than in untreated HCl-injured animals (P <0.01 for both).

### Measurement of the Inflammatory Response

BAL levels of TNF-α (0.09 [0.06–19.00] ng.L^-1^), IL-6 (0.33 [0.18–0.45] ng.L^-1^), IL-1β (0.45 [0.23–0.71] ng.L^-1^) and IL–18 (0.4 [0.2-0.7] ng.L^-1^) were significantly higher in injured animals than in uninjured controls (0.02 [0.01–0.03] ng.L^-1^, 0.02 [0.01–0.03] ng.L^-1^, 0.14 [0.06–0.21] ng.L^-1^ and 0.10 [0.06–0.20] ng.L^-1^; P <10^-4^, P <10^-3^, P <0.01 and P <10^-3^, respectively). By contrast, BAL levels of cytokines were similar in the sham group and in injured animals treated with RAP or sRAGE (**Fig. 4**).

**Figure 4.**
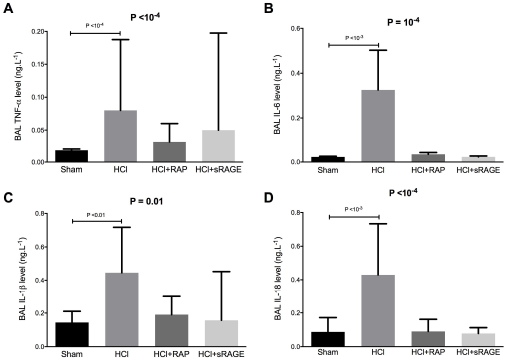
RAGE inhibition decreases alveolar inflammation after acid-induced lung injury. Measurement of bronchoalveolar lavage (BAL) levels of (**A**) tumour necrosis factor (TNF)-α, (**B**) interleukin (IL)-6, (**C**) IL-1β and (**D**) IL-18 in uninjured (Sham), acid-injured (HCl) and acid-injured piglets treated with RAGE antagonist peptide (HCl+RAP) or recombinant sRAGE (HCl+sRAGE) (n = 12 per group). Values are reported as medians and interquartile ranges.

### Histological Evidence of Tissue Injury

Lung injury scores were higher in untreated HCl-injured animals than in sham controls or in HCl-injured animals treated with RAP or sRAGE (**Fig. 5**). When compared to sham animals, alveolar wall thickening and neutrophilic alveolar-interstitial infiltrates were more evident in acid-induced than in non-injured piglets. The ARDS animals treated with RAP or sRAGE had less intense neutrophilic infiltration when compared to untreated acid-injured animals.

**Figure 5.**
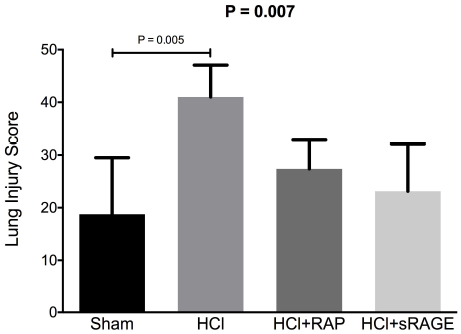
RAGE inhibition decreases histological features of lung injury. Lung injury scores were higher in acid-injured (HCl) than in uninjured piglets (Sham) and acid-injured piglets treated with RAGE antagonist peptide (HCl+RAP) or recombinant sRAGE (HCl+sRAGE) (n = 12 per group). Values are reported as medians and interquartile ranges.

### Other respiratory and hemodynamic variables

The values for tidal volume, PaCO2, mean arterial pressure, cardiac index and serum lactate did not differ between groups over time (**Supplementary Table 3**). However, experimental ARDS was associated with a marked decrease in the compliance of the respiratory system at 1–4 hours after injury, when compared to the baseline, the absence of injury or treatment with RAP and sRAGE. Conversely, the inspiratory plateau and driving pressures were increased after acid injury at 1–4 hours after injury, when compared to the baseline, the absence of injury or treatment with RAP and sRAGE.

### Lung Expression of Alveolar Epithelial Channels

Acid injury was associated with down-regulation of lung mRNA expression of epithelial channels α1-ENaC channel, α1-Na,K-ATPase and AQP-5 (**Fig. 6**). Treatment with either RAP or sRAGE restored lung mRNA expression of AQP-5, whereas treatment with RAP restored lung mRNA expression of α1-Na,K-ATPase. Treatment with RAP or sRAGE had a moderate impact on lung mRNA expression of α1-ENaC that did not reached statistical significance in post-hoc, between-group analysis.

**Figure 6.**
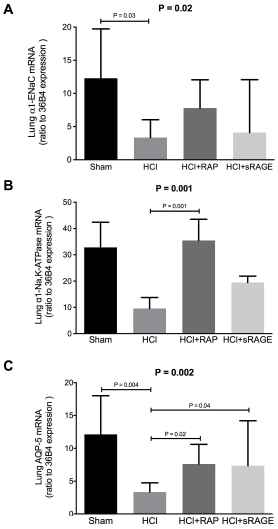
RAGE inhibition restored lung expression of epithelial sodium channel (α1-ENaC), α1-Na,K-ATPase and aquaporin (AQP)-5 after acid-induced lung injury. The gene expression levels of α1-ENaC (NCBI Reference Sequence, NM_213758.2), α1-Na,K-ATPase (NM_214249.1), AQP-5 (NM_001110424.1) and 36B4 (a housekeeping gene) were assessed using the quantitative real-time reverse transcription polymerase chain reaction in uninjured (Sham), acid-injured (HCl) and acid-injured piglets treated with RAGE antagonist peptide (HCl+RAP) or recombinant sRAGE (HCl+sRAGE) (n = 12 per group). Threshold levels of mRNA expression (ΔΔCt) were normalised to the housekeeping gene 36B4 and the values represent the mean of triplicate samples ± SD. Data are representative of three independent observations.

## DISCUSSION

The main goal of this study was to determine the impact of a RAGE inhibition strategy, based either on sRAGE or RAP administration, in a piglet model of HCl-induced ARDS [12,24]. Here, we demonstrated that both sRAGE and RAP had similar beneficial effects on the symptoms of experimental ARDS, including restoration of AFC, improvement in oxygenation, and attenuation of histological lung injury, alveolar-capillary permeability and inflammation. In addition, we proposed that RAGE inhibition might improve AFC in experimental ARDS through restored lung expression of α1-Na,K-ATPase, AQP-5 and, to a lesser extent, α1-ENaC.

Growing preclinical evidence indicates that RAGE modulation might reduce lung injury, although some uncertainty persists [34]. To date, the use of sRAGE as a decoy receptor and the use of anti-RAGE antibody as a direct antagonist have shown beneficial effects on lung injury or sepsis. Indeed, RAGE^-/-^ mice had a survival advantage following cecal ligation puncture (CLP) when compared with wild-type mice [35], and treatment with an anti-RAGE monoclonal antibody also decreased mortality in septic wild-type mice when compared to controls, even when treatment was administered 24 h after CLP. An anti-RAGE monoclonal antibody also improved survival in mice even when the treatment was given 6 h after intratracheal infection with *Streptococcus pneumoniae* [18]. An intraperitoneal sRAGE treatment reduced neutrophil infiltration, lung permeability, proinflammatory cytokine production, NF-κB activation and the number of apoptotic cells in intratracheally LPS-challenged mice [19]. However, another study showed that neither intratracheal nor intraperitoneal sRAGE treatment affected LPS-induced or *Escherichia coli*-induced acute pulmonary inflammation [36]. These conflicting findings on sRAGE may reflect the complexity of RAGE signalling occurring during lung injury [37,38]. Indeed, the RAGE pathway could be considered a double-edged sword, playing important roles both in tissue injury and in resolving the pathogenesis of an offending insult [37].

The role of the balance between RAGE circulating isoforms and ligands in the regulation of RAGE signalling remains poorly investigated under physiological conditions and in specific diseases [38–40]. For example, the use of a model of ventilator-induced lung injury (VILI) using RAGE^-/-^ mice confirmed the contribution of RAGE activation to inflammatory cell influx into the alveolar compartment, but not to other VILI parameters, such as indices of alveolar barrier dysfunction or BAL IL-6, IL-1β and keratinocyte-derived chemokine [41]. By contrast, the use of a two-hit model of VILI and inhaled LPS demonstrated that RAGE^-/-^ mice had elevated cytokine and BAL chemokine levels and that RAGE deficiency did not affect the lung wet-to-dry ratio, total protein level or cell influx [41]. Moreover, administration of sRAGE to RAGE^-/-^ mice attenuated the production of inflammatory mediators, probably because sRAGE can scavenge ligands that have the potential to activate other pattern-recognition receptors such as toll-like receptor 4 [42]. To our knowledge, however, our study is the first to report beneficial effects of S100P-derived RAP administration in experimental ARDS, a finding that agrees with the results from a previous study of HMGB1-derived RAP in lung-injured mice [43].

The current study confirms our own recent results that support the benefits of using sRAGE or an anti-RAGE antibody to restore AFC and attenuate major features of lung injury in a mouse model of acid-induced ARDS [20]. Our current findings also provide further insights into mechanisms by which manipulating the RAGE pathway might counteract ARDS by AFC restoration and lung expression of lung epithelial channels α1-Na,K-ATPase, AQP-5 and, to a lesser extent, α1-ENaC. RAGE activation has been reported to stimulate ENaC activity and lung fluid clearance in uninjured mice via advanced glycation end-products [44]. Conversely, RAGE inhibition was associated with restored AFC and lung AQP-5 expression in acid-injured mice, when compared to controls [20]. However, the precise pathways linking RAGE modulation and active transepithelial fluid transport through the regulation of epithelial barrier integrity and channel activity are not yet established.

Confirming the potential of RAGE modulation in a large animal model is a mandatory step prior to translation of ARDS treatment to the clinical setting. ARDS remains a syndrome that still lacks effective pharmacological therapies, but many issues require resolution before initiating the testing of RAGE inhibition strategies in human patients with ARDS. One issue is that RAGE inhibitors under development can take many forms, including derivatives of sRAGE that may act as decoy molecules, derivatives of RAGE ligands that block membrane RAGE, protein-protein interaction inhibitors, ligand release inhibitors and ligand inactivators [45–48]. Consequently, the choice of a specific agent (or of a combination of agents) deserves full comparative investigations. A second issue is that the intact function of the RAGE pathway may be crucial for antibacterial defence, as RAGE^-/-^ mice showed enhanced bacterial growth, increased bacterial dissemination and more severe inflammation in a model of bacterial peritonitis [49]. However, contradictory findings have been published and other studies have shown that RAGE inhibition, elicited either through genetic deletion or through the use of anti-RAGE or anti-HMGB1 antibodies, was associated with unchanged or decreased bacterial dissemination [35,50,51]. Indeed, although trials in patients with mild Alzheimer’s disease did not support clinical efficacy of azeliragon (TTP448, vTv Therapeutics, High Point, NC, USA), an inhibitor of RAGE-amyloid β protein interactions, no obvious safety issues were associated with its use (5 mg orally once daily for 18 months) in human patients [52–54]. Future research should therefore investigate the extent to which our current findings might translate to the treatment of critically ill patients with ARDS in terms of the timing, dosing and methods of administration of RAGE inhibition candidates, with a focus on their efficacy and safety profiles.

Our study has some limitations. One limitation is that we mainly focused on the major criteria of experimental ARDS, including AFC measurements [33], and the assessment of animals was limited to four hours after injury. Therefore, the extrapulmonary and longer-term effects of RAGE inhibition in this model remain unknown. A second limitation is that we suggested restored AFC and lung expression of epithelial channels as potential mechanisms for the beneficial effects of RAGE inhibition, but we did not precisely characterise the specific pathways by which RAP and sRAGE might alleviate lung injury in ARDS, and this deserves further investigation. Furthermore, our findings may hold true only for acid-induced ARDS, so additional validation is warranted in other settings, such as pulmonary and extrapulmonary sepsis (the most frequent cause of ARDS), prior to considering clinical translation into more complex critical care scenarios that frequently combine multiple organ failure.

In conclusion, a RAGE inhibition strategy, using either recombinant sRAGE or RAP, was associated with restored AFC and attenuated lung injury in a translational piglet model of acid-induced ARDS. These results, which reinforce those from previously published preclinical studies in smaller animals, might represent an important step towards future clinical translation of this treatment strategy, although further investigations are needed to confirm the safety of modulating RAGE in patients with ARDS.

## Author Contributions

Jules Audard, Vincent Sapin, Jean-Michel Constantin, and Matthieu Jabaudon conceived and designed the study. Jules Audard, Thomas Godet, Raiko Blondonnet, Jean-Baptiste Joffredo, and Matthieu Jabaudon performed the experiments. Jules Audard, Raiko Blondonnet, Bertille Paquette, Corinne Belville, Marlyne Lavergne, Christelle Gross, and Matthieu Jabaudon collected and analysed the data. Jules Audard, Thomas Godet, Justine Pasteur, Damien Bouvier, Loic Blanchon, Bruno Pereira, and Matthieu Jabaudon interpreted the data. Vincent Sapin, Jean-Michel Constantin and Matthieu Jabaudon acquired the funding and supervised the study. Jules Audard and Matthieu Jabaudon wrote the paper, all other authors reviewed and edited the manuscript prior to submission.

## Supplementary Material

The ARRIVE Guidelines Checklist

Animal Research: Reporting In Vivo Experiments

Carol Kilkenny^1^, William J Browne^2^, Innes C Cuthill^3^, Michael Emerson^4^ and Douglas G Altman^5^

^1^The National Centre for the Replacement, Refinement and Reduction of Animals in Research, London, UK, ^^2^^School of Veterinary Science, University of Bristol, Bristol, UK, ^^3^^School of Biological Sciences, University of Bristol, Bristol, UK, ^^4^^National Heart and Lung Institute, Imperial College London, UK, ^^5^^Centre for Statistics in Medicine, University of Oxford, Oxford, UK.

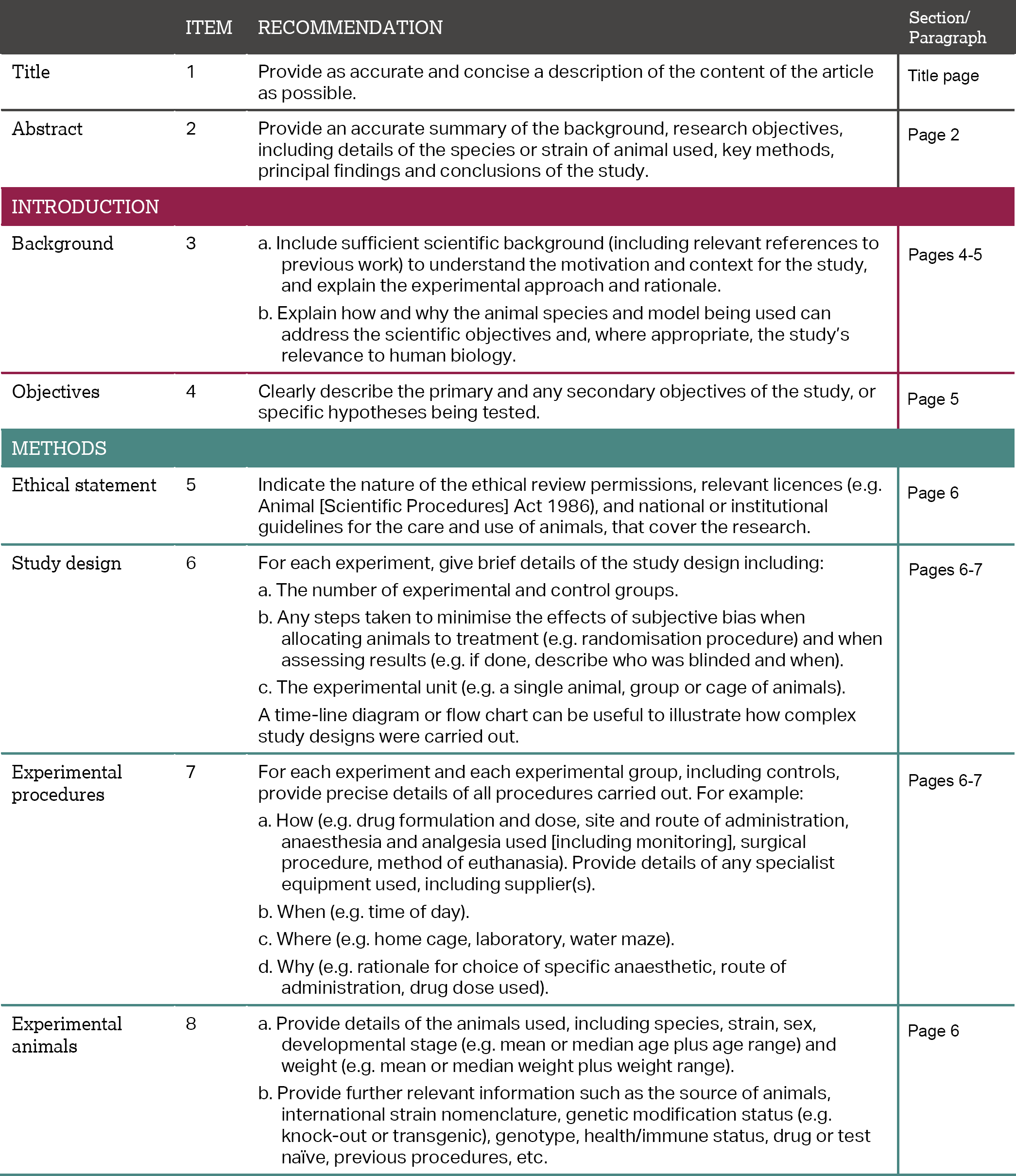
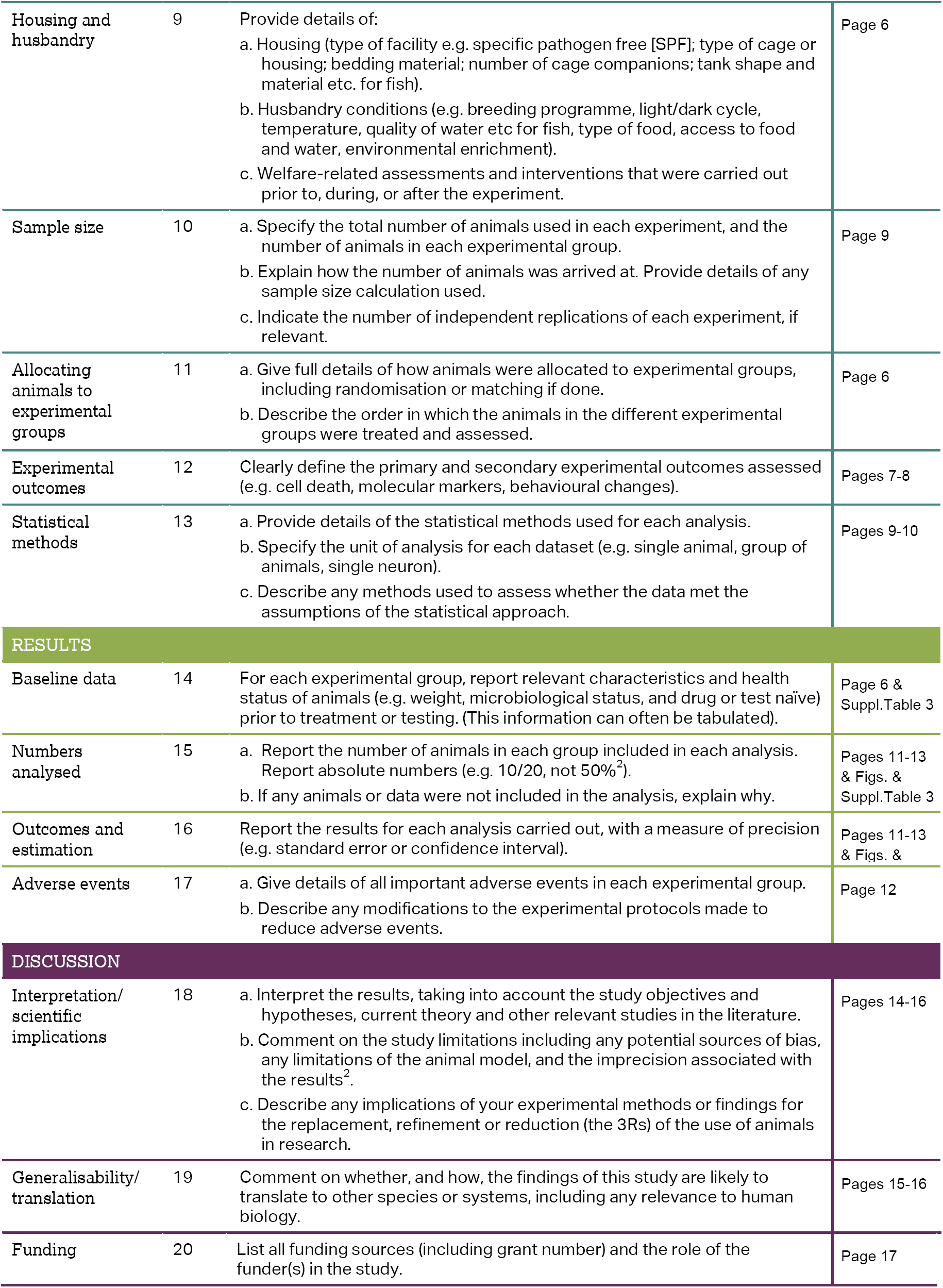

**Table 1.** Lung Histology Injury Scoring System (adapted from Matute-Bello et al. [33]).

| Parameter | 0 | 1 | 2 |
| --- | --- | --- | --- |
| i. Neutrophils in the alveolar space | None | 1–5 | >5 |
| ii. Neutrophils in the interstitial space | None | 1–5 | >5 |
| iii. Hyaline membranes | None | 1 | >1 |
| iv. Proteinous debris filling the airspaces | None | 1 | >1 |
| v. Alveolar septal thickening | None | 2×–4× | >4× |

**Table 2.**
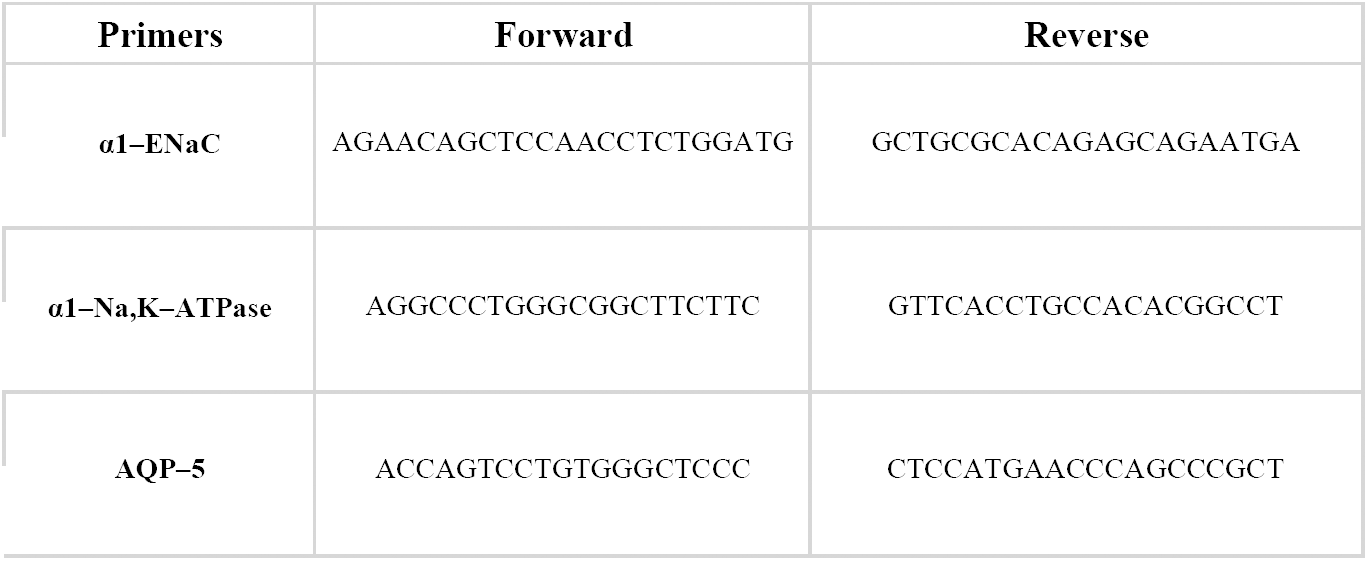
Gene-Specific Primer Sequences Used in the Study.

**Table 3.**
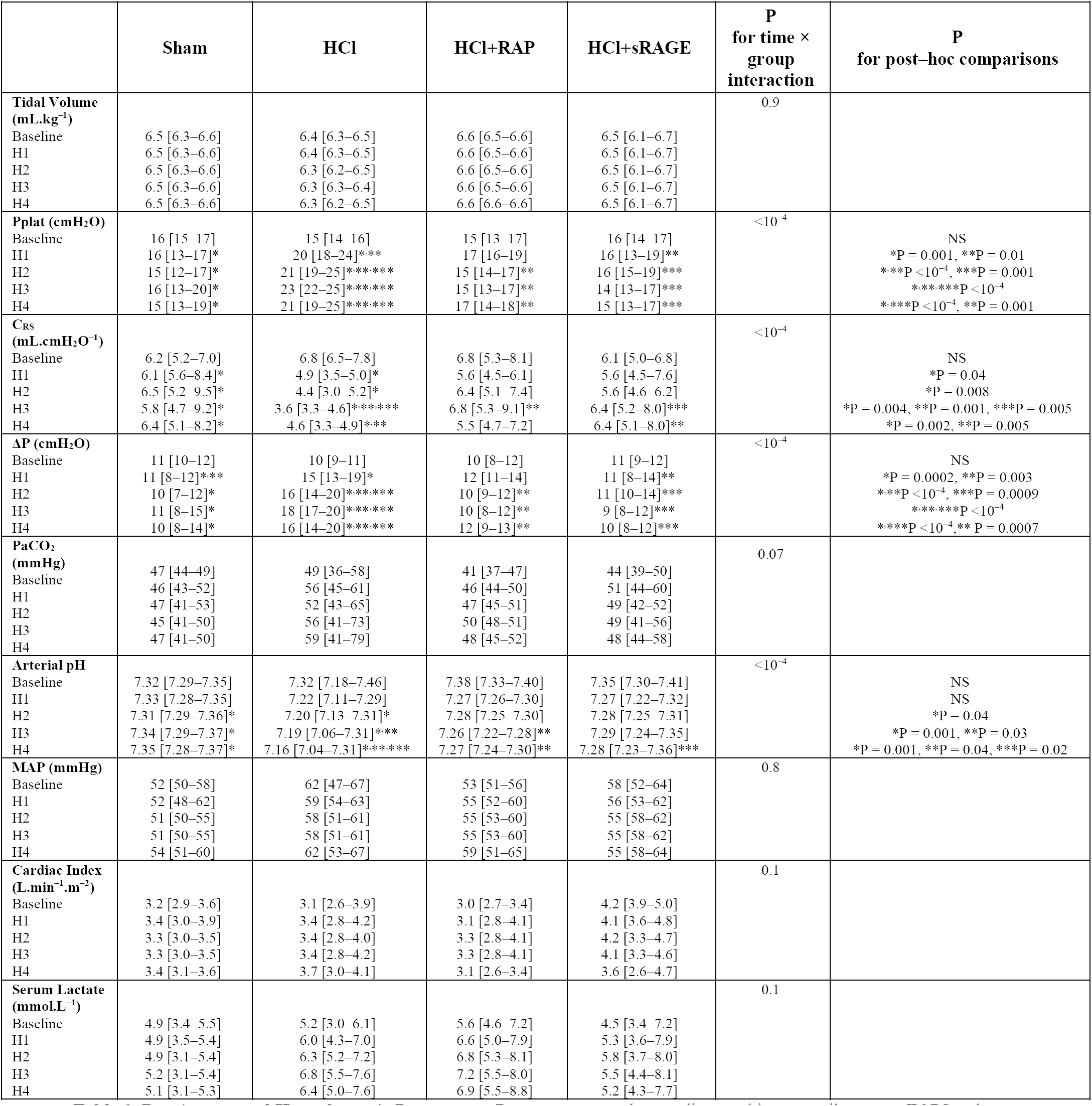
Respiratory and Hemodynamic Parameters. Data are presented as medians and interquartile ranges [IQR] and are analysed with two-way repeated-measurement analysis of variance. When significant, Mann-Whitney test (nonparametric data) was used for post–hoc comparisons between groups at each time point. *Pplat: inspiratory plateau pressure. C*_*RS*_: *compliance of the respiratory system. ΔP: driving pressure. PaCO*_*2*_*: arterial carbon dioxide tension. MAP: mean arterial pressure. NS: non-significant.*

